# Loss of the Parkinson’s disease-associated protein DJ-1 impacts dopamine metabolism in astrocytes

**DOI:** 10.1101/2025.11.02.685006

**Authors:** Annika Wagener, Naiyareen Mayeen, Stephan A. Müller, Sarah K. Tschirner, Brigitte Nuscher, Pauline Mencke, Stefan F. Lichtenthaler, Ibrahim Boussaad, Rejko Krüger, Lena F. Burbulla

## Abstract

The selective loss of dopaminergic neurons in the substantia nigra is a hallmark of Parkinson’s disease (PD), yet the contribution of glial cells to this vulnerability is not fully understood. Studies in rodent models suggest that astrocytes can take up and metabolize dopamine (DA), potentially protecting neurons by detoxifying reactive DA metabolites via glutathione S-transferase mu 2 (GSTM2) release. However, these mechanisms remain underexplored in human systems, particularly in the context of PD. Here, we used CRISPR-engineered iPSC-derived human astrocytes with a PD-linked DJ-1 mutation and isogenic controls to investigate astrocytic DA metabolism. Upon DA exposure, control astrocytes upregulated quinone-reducing enzymes NAD(P)H quinone dehydrogenase 1 (NQO1) and GSTM2, whereas DJ-1 mutant astrocytes failed to adaptively respond. In addition, only control astrocytes presented with increased DA quinone products upon DA exposure, not DJ-1 mutants. These results demonstrate astrocytic DA handling being disrupted in DJ-1-linked PD, implicating astroglial dysfunction as an important contributor to PD pathogenesis and potential target for therapeutic intervention.

## Introduction

The selective degeneration of dopaminergic neurons in the substantia nigra (SN) and the accumulation of α-synuclein in Lewy bodies are hallmarks of Parkinson’s disease (PD), the second most common neurodegenerative disorder. Understanding why dopaminergic neurons are specifically vulnerable in PD remains difficult, despite genetic studies implicating converging pathways such as mitophagy, lysosomal function, and oxidative stress (*1*). Among these, mutations in *PARK7,* encoding the redox-sensitive protein DJ-1, are known to affect mitochondrial health and oxidative stress regulation in familial and sporadic forms of PD (*2*). While the oxidation of DJ-1 itself activates its antioxidant function, over-oxidation of DJ-1 results in a partially dysfunctional form, as observed in sporadic PD patient brains (*3*). Mutations in DJ-1 impair its chaperone function and impact its ability to protect cells from oxidative stress, however, the exact mechanisms linking DJ-1 to PD pathogenesis are still undefined (*4*). Using iPSC-derived midbrain dopaminergic patient neurons, we previously showed that loss of DJ-1 triggers a time-dependent pathologic cascade in which mitochondrial oxidative stress facilitates the accumulation of oxidized derivates of dopamine (DA), i.e. DA quinone products, itself mediating lysosomal dysfunction and α-synuclein accumulation (*5*).

Neuronal metabolism of DA is critical for maintaining cellular homeostasis due to its high oxidative liability. Under physiological conditions, DA released into the synaptic cleft is rapidly cleared by DA transporter (DAT) and either re-sequestered into synaptic vesicles by vesicular monoamine transporter 2 (VMAT2) or enzymatically degraded in the cytosol by monoamine oxidase (MAO) or catechol-O-methyltransferase (COMT). One intermediate of this catabolic degradation of DA is the highly reactive intermediate 3,4-dihydroxyphenylacetaldehyde (DOPAL), which is subsequently converted by aldehyde dehydrogenase into the non-toxic derivate 3,4-dihydroxyphenylacetic acid (DOPAC). Both free cytosolic DA and DOPAL are prone to further oxidation (*6*) and readily form reactive quinones, such as aminochrome, which can inactivate proteins, promote α-synuclein aggregation, trigger inflammation, and impair mitochondrial complex I function (*7–9*).

Therefore, cells rely heavily on antioxidant and detoxification pathways to counter the harmful effects of DA-derived quinones. One such pivotal mechanism involves glutathione S-transferase mu 2 (GSTM2), which catalyses the conjugation of aminochrome with glutathione to yield non-toxic derivatives, thus protecting against quinone-induced cellular damage (*10, 11*). NAD(P)H quinone dehydrogenase 1 (NQO1) (also known as DT-diaphorase) complements this process by reducing quinones to hydroquinones, thereby preventing reactive oxygen species generation (*12*). Both enzymes protect DA neurons against oxidative stress *in vitro* and *in vivo* (*13–15*).

Although much of PD research has been focused on neurons, the role of astrocytes as important contributors to DA neuron degeneration is receiving growing attention (*16*). Astrocytes sustain neuronal health by providing metabolic and structural support, maintaining the blood-brain-barrier and neurotrophic signalling (*17*). However, mutations in PD-related proteins have been shown to disrupt critical functions in astrocytes including autophagy, leading to α-synuclein and protein accumulation (*18–21*) as well as increased inflammation, mitochondrial dysfunction, altered ER stress, impaired trophic support, proliferation defects and calcium dysregulation (*22–24*).

Given the central role of DA metabolism in PD pathogenesis, it is increasingly recognized that astrocytes may play a direct and active role in shaping the neuronal microenvironment through their own handling of DA. However, very little is known about how astrocytes metabolize DA and by doing so potentially protect neurons against DA-derived quinone stress. Initial evidence hints towards astrocytes taking up excess DA (*25–28*) through one of their multiple DA uptake systems including DAT, plasma membrane monoamine transporter (PMAT), organic cation transporter 3 (OCT3), and Norepinephrine transporter (NET) (*25, 29, 30*). Astrocytes also contain enzymes for DA packaging and metabolism such as VMAT2 for vesicular storage and COMT and MAO for DA breakdown (*31, 32*). Yet, the physiological relevance and regulation of these DA handling pathways in astrocytes, particularly under pathological conditions like PD, remain unclear.

Importantly, previous studies suggest a potential protective role of astrocytes helping neurons to neutralize the toxicity of DA-derived quinones, particularly the most neurotoxic quinone aminochrome, in dopaminergic neurons, by expressing NQO1 and GSTM2 (*26, 33, 34*). Interestingly, aminochrome has been suggested to drive downstream expression of GSTM2 (*35*). Moreover, GSTM2 is uniquely expressed in human astrocytes and suggested to be secreted in exosomes, which are then taken up by neighbouring dopaminergic neurons, enhancing neuronal resistance to DA quinone and aminochrome toxicity (*36*). However, much of this evidence comes from rodent models or astrocytoma lines and disease-relevant human model system are needed to decipher whether and how human astrocytes manage DA uptake and metabolism, or contribute to quinone detoxification and neuronal protection.

In this study, we investigated iPSC-derived human astrocytes deficient of functional DJ-1 compared to isogenic controls. Interestingly, we uncovered a specific impairment in the metabolic response of DJ-1-deficient astrocytes to DA while healthy control astrocytes responded adequately. While healthy astrocytes robustly upregulated NQO1, GSTM2 and DA quinones in response to DA exposure, DJ-1-deficient astrocytes lack these physiologically relevant reactions. We therefore conclude that DA metabolic and detoxification mechanisms are disrupted in DJ-1-deficient astrocytes. Our findings uncover the loss of the PD-associated protein DJ-1 to impact DA metabolism in astrocytes reducing their neuron-supporting function, intriguingly relevant to PD pathology. Studying how astrocytes could aid vulnerable dopaminergic neurons could be a crucial target for future therapeutic interventions.

## Results

### Loss of DJ-1 impairs mitochondrial respiration and enhances stress signatures in human iPSC-derived astrocytes

To investigate astrocyte-specific mechanisms in PD, we generated astrocytes from a healthy individual (control) and its CRISPR-engineered iPSC line carrying the PD-linked *PARK7*/DJ-1 mutation c.192G>C (DJ-1 mut) previously characterized by Boussaad and co-workers (*37*) using an established protocol (*38*). Immunoblotting analysis demonstrated that both lines exhibited robust expression of the astrocyte marker glial fibrillary acidic protein (GFAP), while lacking expression of the neuronal marker microtubule-associated protein 2 (MAP2) or the marker for midbrain dopaminergic neurons tyrosine hydroxylase (TH), displaying the efficient and selective astrocytic differentiation (Fig. 1A). The absence of DJ-1 protein in the mutant line was confirmed, whereas the control astrocytes displayed strong DJ-1 expression (Fig. 1A). Furthermore, flow cytometry analysis showed GFAP positivity, but absence of MAP2 in differentiated astrocyte cultures (Fig 1B). Additionally, immunocytochemical analysis indicated robust expression of the canonical astrocyte markers GFAP, S100β, Sox9, Vimentin and EAAT2 in both astrocytic lines while lacking the neuronal marker MAP2, consistent with a glial identity (Fig. 1C). To assess the functional competence of the differentiated cells, we evaluated their ability to uptake glutamate, a key physiological property of astrocytes critical for synaptic homeostasis (*39*). Differentiated astrocytes demonstrated efficient glutamate uptake, confirming characteristics of mature, functional astrocytes (Fig. 1D).

**Figure 1:**
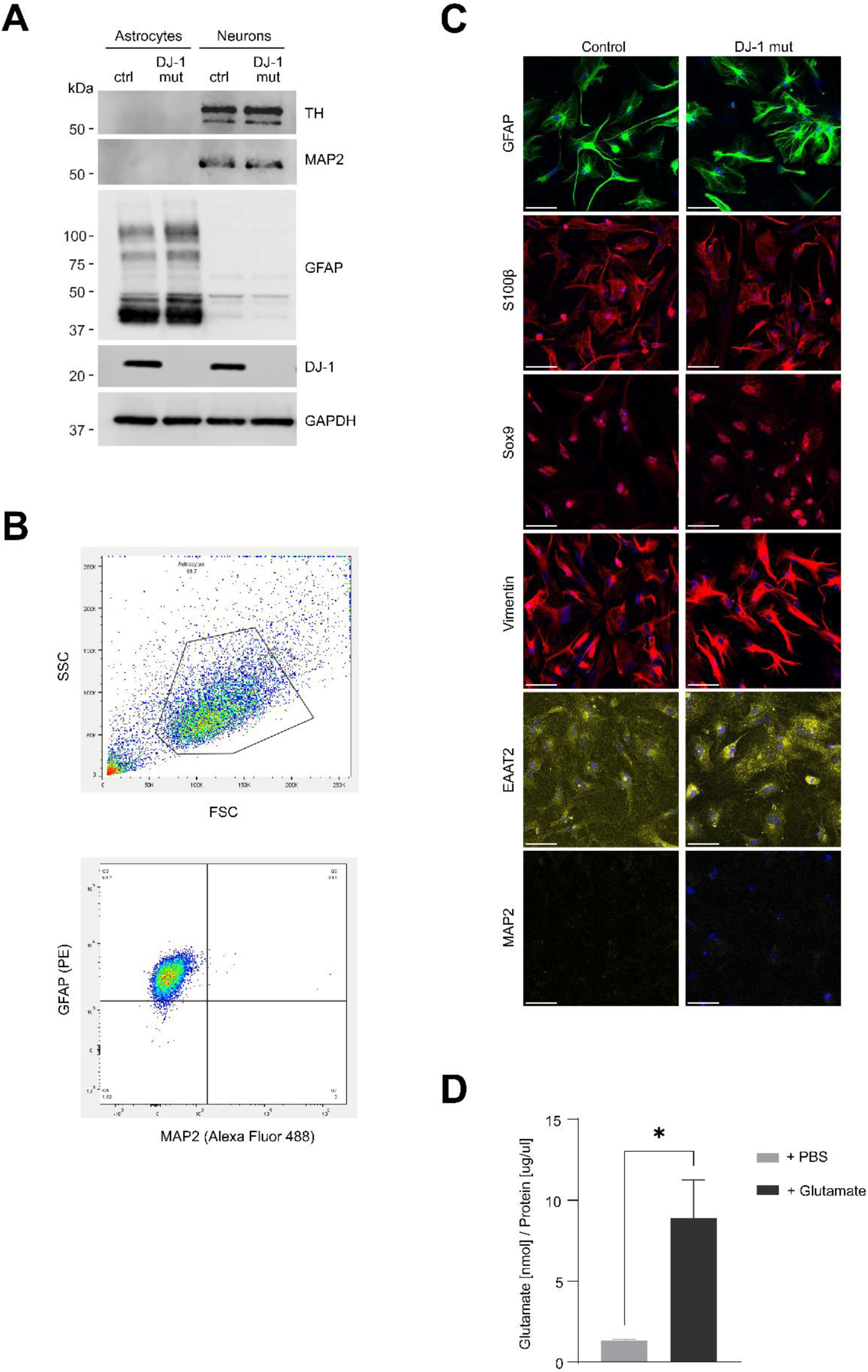
Functional profiling of iPSC-derived astrocytes. (**A**) Immunoblot analysis of the astrocyte marker GFAP, neuronal marker MAP2, midbrain neuron marker TH, and PD-associated DJ-1 in CRISPR-engineered DJ-1-deficient (DJ-1 mut) astrocytes compared to isogenic control. GAPDH was used as loading control. (**B**) Representative image of flow cytometry analysis shows highly pure astrocyte cultures positive for GFAP and negative for MAP2. (**C**) Immunocytochemistry analysis for canonical astrocyte markers GFAP, S100β, Sox9, Vimentin and EAAT2, and the neuronal marker MAP2 in control and DJ-1-deficient astrocytes. Staining shows DAPI as nuclear stain. Scale bar, 50 µM. (**D**) Glutamate uptake assay showing efficient glutamate uptake by control astrocytes (n=4).

Next, mass spectrometry-based proteomics was used as an unbiased approach to identify proteins and pathways associated with disease. We identified 702 proteins to be upregulated and 1282 proteins to be downregulated in DJ-1-deficient astrocytes (Student’s t-test p < 0.05 and log2 fold change larger than 0.5 or lower than −0.5) compared to healthy control astrocytes (Fig. 2A; Supplementary Table). Proteomic analysis of DJ-1-deficient astrocytes revealed significant alterations in proteins associated with mitochondrial function and oxidative stress regulation. Notably, downregulation of FLCN, GLRX2 and SFXN4 suggests compromised mitochondrial redox homeostasis, complex I formation and calcium handling (*40–42*). Concurrent upregulation of stress-response proteins such as HMGB1, HMGB2 and RNASEH2A indicating a cellular response to DNA damage and repair, as well as pro-inflammatory mechanisms, further supports an enhanced oxidative stress state (*43, 44*) (Fig. 2A). Of note, proteomics analysis also indicated the upregulation of COMT in DJ-1-deficient astrocytes, a protein preferentially expressed in glia cells and mainly responsible for the enzymatic degradation of DA.

**Figure 2.**
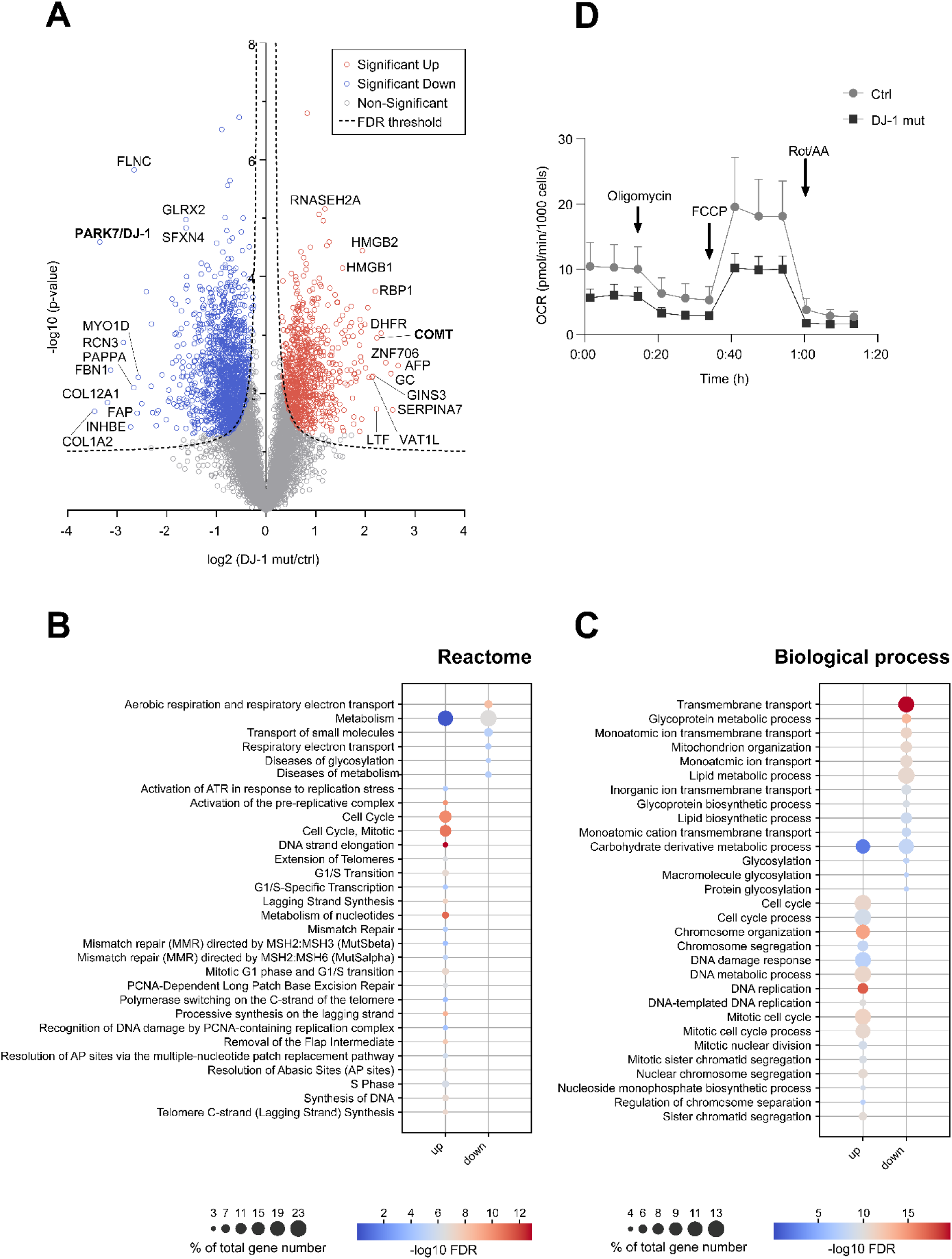
Proteomic analysis reveals mitochondrial deficits in astrocytes upon DJ-1 loss. (**A**) Volcano plot comparing DJ-1-deficient (DJ-1 mut) versus control (WT) astrocytes. The -log10-transformed Student’s t-test p-values of each protein are plotted against their log2-transformed fold change of protein label free quantification intensities. Hyperbolic curves represent a permutation-based FDR threshold (0.05, s_0_=0.1). Proteins are displayed as follows: FDR-significant up (red) and down (blue) p < 0.05 and non-significant (grey) (n = 5 per group). (**B-C**) Enrichment analysis of the DJ-1-deficient astrocyte proteome for (B) Reactome and (C) gene ontology biological processes (GOTERM_BP_DIRECT) using the web tool DAVID (v 6.8) (*62*). Differentially expressed proteins (t-test p-value < 0.05 and log2 fold change > 0.5 or < −0.5) were compared to the background of all proteins detected under both conditions. In the dot plots, dot size represents the total gene number of a term in percent, while colour indicates the -log10-transformed FDR. The top 30 enriched terms are shown. (**D**) Mitochondrial stress test trace of oxygen consumption rate (OCR) following sequential addition of oligomycin, FCCP, rotenone, and antimycin A in DJ-1-deficient compared to control astrocytes (n=6), p=0.0004.

Additionally, we performed an enrichment analysis for Reactome pathway (Fig. 2B), as well as Gene Ontology (GO) biological process (Fig. 2C), molecular function (Fig. S1A) and cellular component (Fig. S1B) of these differentially expressed proteins to further investigate their associated signaling pathways (Supplementary Table). We ranked the top 30 terms by significance and discovered a prominent enrichment of downregulated proteins for the terms “Mitochondrion organization”, “Mitochondrion”, “Aerobic respiration and respiratory electron transport” and “Respiratory electron transport” associated with biological processes, cellular compartment or Reactome pathway in DJ-1-deficient astrocytes. On the other hand, the analysis revealed a prominent enrichment of upregulated proteins for terms “DNA replication”, “DNA damage response” and “Cell Cycle” associated with biological processes or Reactome pathway, respectively, in DJ-1-deficient astrocytes. These results indicate metabolic disturbances and specifically hint towards major mitochondrial dysfunction and cellular stress response in mutant astrocytes, possibly compromising their proper function in maintaining neuronal homeostasis.

To validate mitochondrial function in control and DJ-1 mutant astrocytes, we performed the mitochondrial stress test assay using the Seahorse XF Analyzer. This approach monitors key bioenergetic parameters in real time by measuring oxygen consumption rate (OCR) after sequential addition of oligomycin, FCCP, and a mixture of rotenone and antimycin A (Rot/AA) (Fig. 2D). Notably, DJ-1-deficient astrocytes exhibited a reduced OCR compared to isogenic controls. In summary, loss of DJ-1 in human iPSC-derived astrocytes results in marked mitochondrial impairment and triggers broad cellular stress pathways, revealing a vulnerability that may compromise astrocyte-mediated neuronal support in PD.

### DJ-1 deficiency impairs dopamine metabolism in human astrocytes

To assess astrocytic DA metabolism, control and DJ-1-deficient iPSC-derived astrocytes were treated with 60 μM DA or PBS as vehicle and collected for High-Performance Liquid Chromatography (HPLC) analysis to quantify DA and its primary metabolite 3-dihydroxyphenylacetic acid (DOPAC). Representative chromatograms show that, as expected, PBS-treated samples only presented the internal standard 3,4-dihydroxybenzylamine (DHBA) and background peaks at the beginning of the HPLC measurement (Fig. 3A, top), whereas astrocytes exposed to DA displayed prominent peaks corresponding to both DA and DOPAC, indicating uptake and metabolism of exogenous DA (Fig. 3A, bottom). Subsequent HPLC analysis for quantification of intracellular DA content revealed an increase in both control and DJ-1-deficient astrocytes following DA exposure compared to PBS, confirming that both cell lines are capable of efficient DA uptake (Fig. 3B). Notably, there was no significant difference in overall DA content after uptake between control and DJ-1-deficient astrocytes, indicating that loss of DJ-1 does not impair DA uptake capacity. However, analysis of the DOPAC to DA ratio demonstrated a reduced efficiency of DA metabolism into its non-toxic metabolite under DJ-1-deficient conditions in human astrocytes (Fig. 3C). This suggests that while DJ-1-deficient astrocytes can take up DA, their ability to metabolize it to DOPAC is compromised, highlighting a defect in DA metabolic processing in the absence of DJ-1.

**Figure 3.**
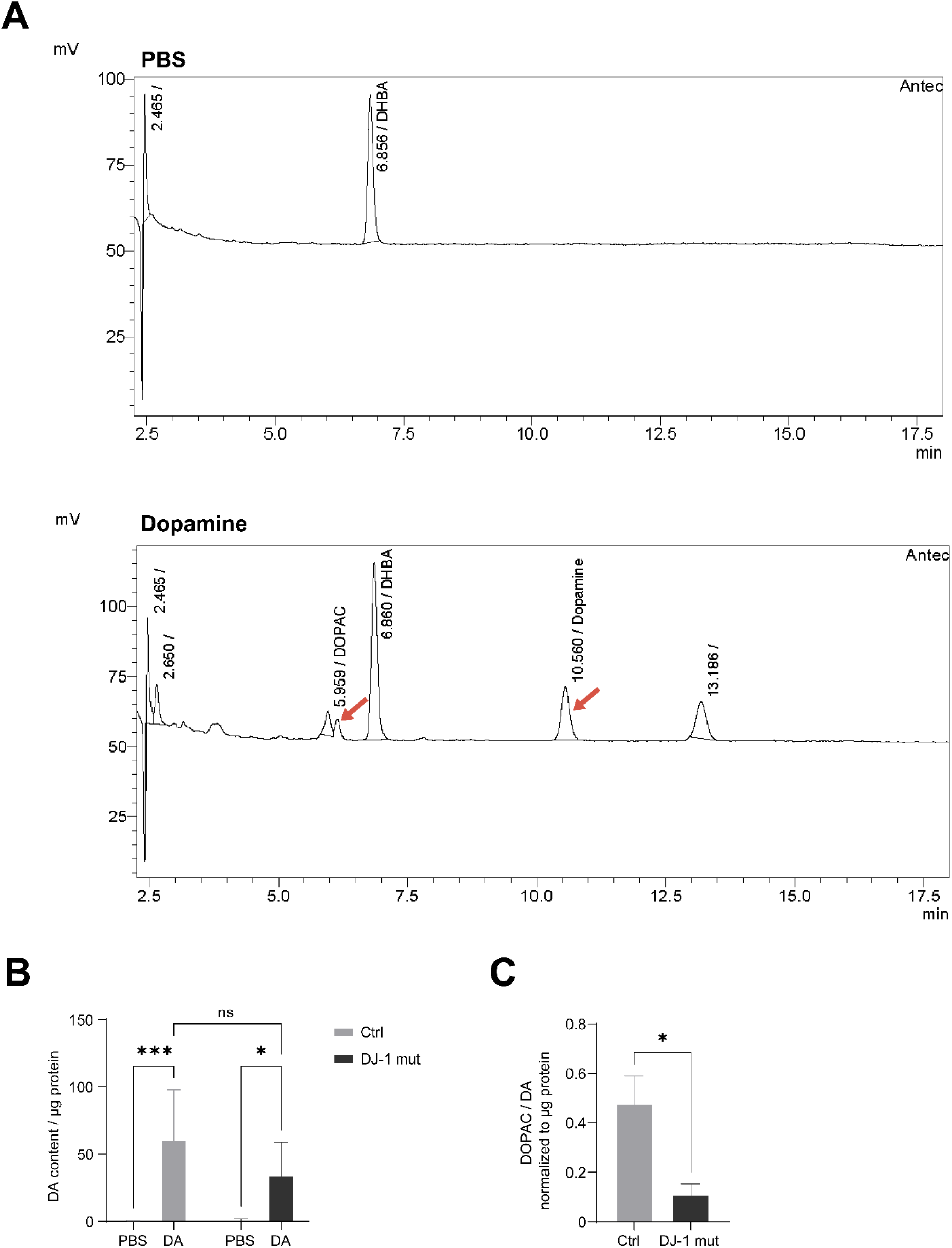
Loss of DJ-1 reduces metabolic conversion of dopamine to DOPAC in iPSC-derived astrocytes following dopamine uptake. (**A**) Representative HPLC chromatograms of cell extracts from PBS (top traces) or 60 μM DA-treated (bottom traces) astrocytes. Only internal standard (DHBA) is detected in PBS, while DA-treated samples display peaks for DA and DOPAC, indicated by red arrows. (**B**) Quantification of intracellular DA content in control and DJ-1-deficient astrocytes after DA treatment (n=5). (**C**) DOPAC to DA ratio in control and DJ-1-deficient astrocytes (n=5).

### Loss of DJ-1 hinders activation of antioxidant pathways in human astrocytes upon dopamine exposure

Building on the observed impairment of conversion of DA to DOPAC in DJ-1-deficient astrocytes, we investigated quinone-reducing enzymes, key players of detoxification of DA and toxic DA derivates. First, control and DJ-1 mutant astrocytes were treated with DA or vehicle, and levels of the vesicular monoamine transporter 2 (VMAT2) – important for the sequestration of DA into vesicles – were assessed (Fig. 4A). Exposure to DA had no effect on VMAT2 expression in control or DJ-1-deficient astrocytes.

**Figure 4.**
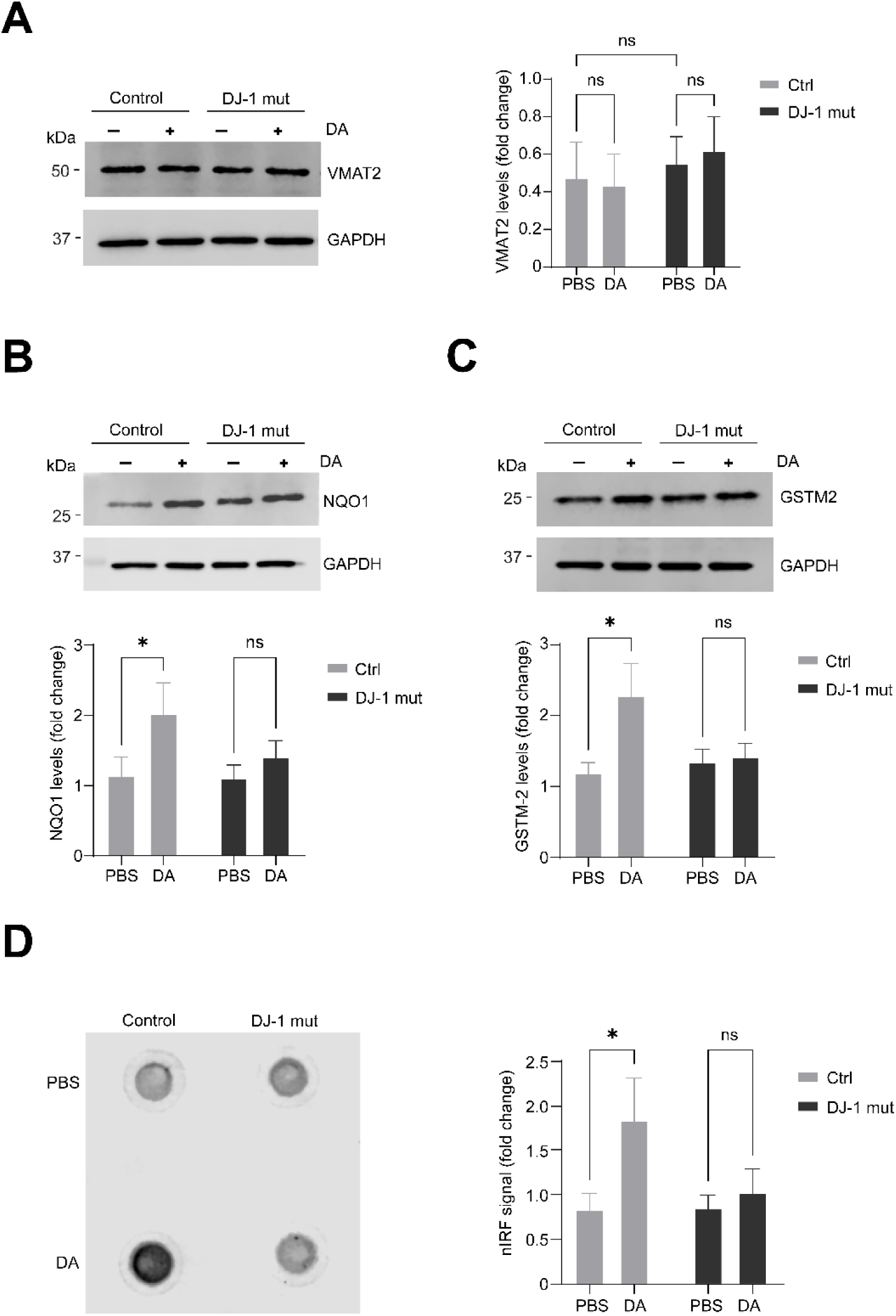
DJ-1 deficiency disrupts dopamine-induced antioxidant responses in human iPSC-derived astrocytes. (**A-D**) Representative immunoblot and quantification of (A) VMAT2 protein levels (n=9), (B) NQO1 protein levels (n=7), and (C) GSTM2 protein levels (n=7) in control and DJ-1-deficient astrocytes treated with PBS or DA. GAPDH was used as loading control. (D) Representative nIRF assay showing oxidized DA signal under baseline (PBS) and DA-treated conditions, and quantification (n=8).

Next, we examined the key antioxidant players NAD(P)H:quinone acceptor oxidoreductase 1 (NQO1) and Glutathione S-Transferase Mu 2 (GSTM2), which are both known to detoxify aminochrome. DA treatment significantly increased NQO1 protein levels in control astrocytes, whereas DJ-1-deficient cells failed to upregulate NQO1 in response to DA (Fig. 4B). Consistent with the lack of NQO1 induction in DJ-1-deficient astrocytes, GSTM2 protein abundance also remained unchanged in mutant astrocytes upon DA treatment, while DA exposure led to an elevation of GSTM2 levels in control astrocytes (Fig. 4C). Finally, we quantified DA quinones, including aminochrome, using a near-infrared fluorescence (nIRF) assay (*45*), and observed a significant increase of these species in control astrocytes upon DA exposure (Fig. 4D). As suggested in previous studies aminochrome exposure may be a relevant trigger of GSTM2 expression (*35*). In contrast, DJ-1-deficient astrocytes showed no elevation in DA quinone levels following DA treatment (Fig. 4D).

Collectively, these findings indicate that DJ-1 is essential for triggering a coordinated astrocytic antioxidant response to DA challenge, including the induction of key antioxidant enzymes NQO1 and GSTM2. Loss of DJ-1 disrupts these protective responses. This failure of glial adaptation in DJ-1-deficient astrocytes is hypothesized to increase the vulnerability of neighbouring neurons due to impaired upregulation of astrocytic GSTM2 following DA stimulation, possibly resulting in diminished neuroprotection.

## Discussion

Our study demonstrates that DJ-1 deficiency impairs mitochondrial respiration and disrupts DA metabolic processing in human astrocytes, providing new insights into glial contributions to PD pathogenesis. Using disease modelling strategies, healthy control and DJ-1-deficient astrocytes were differentiated from iPSCs to investigate their mitochondrial and proteomic signature as well as neuro-supportive strategies.

Findings on marked mitochondrial and bioenergetics disturbances in DJ-1-deficient astrocytes are consistent with previous publications implicating a role of DJ-1 in protecting mitochondrial integrity and minimizing oxidative stress in both neurons and glia (*5, 46*). Additional studies highlight the potential of astrocytic mitochondrial impairment to disrupt brain homeostasis and amplify neuronal vulnerability in PD (*16, 47*). Interestingly, astrocytes with DJ-1 deficiency demonstrate a markedly reduced capacity for neuroprotection, an effect mediated by the loss of neuroprotective soluble factors present in conditioned media, including antioxidant molecules and neurotrophic or bioenergetic proteins essential for neuronal resilience against mitochondrial toxins (*48, 49*).

One crucial aspect of astrocyte-neuron communication, is the capacity of astrocytes to uptake and thus regulate neurotransmitters, including DA (*26, 50*). Notably, we show that both control and DJ-1-deficient astrocytes were able to uptake DA efficiently. However, DJ-1-deficient astrocytes displayed reduced conversion of DA to its non-toxic metabolite DOPAC, as reflected by the decreased DOPAC/DA ratio. These results indicate a specific defect in DA metabolic processing rather than uptake. Surprisingly, this effect was paralleled by the accumulation of oxidized DA species in control, but not DJ-1-deficient astrocytes. We hypothesize that the observed upregulation of COMT in DJ-1-deficient astrocytes, detected in our proteomic analysis diminished the amount of cytosolic DA after astrocytic uptake, thus by-passing the generation of aminochrome and failing to activate antioxidant defense mechanisms in astrocytes (*6*). Through which mechanisms the expression of COMT as a major DA degradation enzyme is initiated in a DJ-1 deficiency scenario requires further investigation. However, we hypothesize that as a result of rapid DA degradation, downstream metabolic antioxidant pathways are disrupted and the capacity of DJ-1-deficient astrocytes to aid neurons coping with neuronal DA-induced oxidative stress, such as neuronal aminochromone built-up, may be substantially diminished.

In fact, we observed that the antioxidant enzymes NQO1 and GSTM2 were robustly induced by DA in control astrocytes, but completely failed to be upregulated in DJ-1-deficient astrocytes. Both enzymes are essential for detoxifying DA-derived quinones and reactive intermediates, serving as key elements of the astrocytic antioxidant machinery (*51*). Interestingly, DJ-1 deficiency in a human cancer cell line and mouse embryonic fibroblasts leads to an impairment of NQO1 stability and function, increasing neuronal susceptibility to oxidative damage (*52*). Importantly, emerging evidence indicates that GSTM2 is produced in astrocytes with inducible expression following aminochrome exposure (*35*), and is also secreted and taken up by neighbouring neurons, where it plays a direct neuroprotective role by aiding in the detoxification of neuronal aminochrome (*53*). This mechanism is proposed to be mediated by exosomes, which have been shown to contain GSTM2 in fibroblasts (*36, 54*).

As such, a failure of DJ-1-deficient astrocytes to upregulate (and possibly secrete) GSTM2 in response to DA may further diminish this neuroprotective support. This could contribute to the long-discussed selective vulnerability of midbrain dopaminergic neurons in PD, and at the same time, highlights GSTM2 supplementation or the enhancement of astrocyte-neuron metabolic cooperation as intriguing therapeutic strategies.

Collectively, our results position DJ-1 as a central regulator of astrocytic defence against mitochondrial and dopaminergic stress, orchestrating both basal metabolic function and induced antioxidant gene expression (Fig. 5). The inability of DJ-1-deficient astrocytes to metabolize DA properly and launch a coordinated antioxidant response to DA challenge may heighten their risk of oxidative injury and impair their capacity to protect neurons from oxidative damage. Importantly, these findings extend the focus of DJ-1-related PD pathogenesis from neurons to astrocytes and suggest that restoration of glial bioenergetic and detoxification capacity may offer new therapeutic avenues, including potential strategies aimed at enhancing GSTM2-mediated astrocyte-to-neuron support.

**Figure 5:**
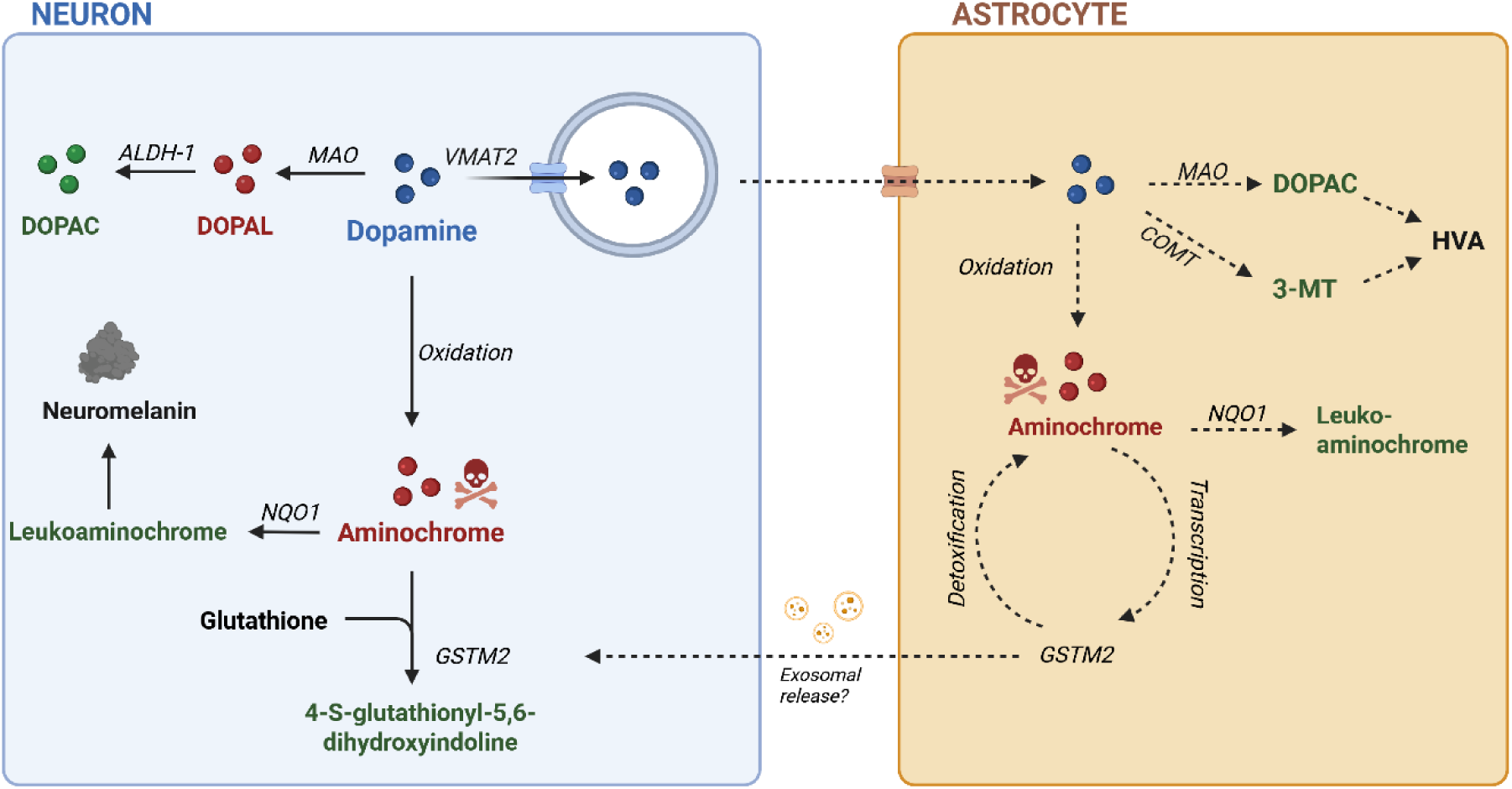
Astrocytic dopamine uptake, metabolism, quinone detoxification and GSTM2-mediated neuroprotection via neuronal uptake. Astrocytes take up extracellular DA, after which DA is metabolized through MAO and COMT to form DOPAC, 3-MT, and HVA. However, a notable fraction of internalized DA undergoes enzymatic oxidation to generate aminochrome. Detoxification of aminochrome is achieved by NQO1, which reduces it to leukoaminochrome, and by GSTM2, which catalyses its conjugation to glutathione, forming non-toxic products and thereby protecting against quinone-induced damage. Moreover, aminochrome stimulates transcriptional upregulation of GSTM2, suggested to be released through exosomes. Astrocyte-derived GSTM2, when taken up by neurons, is considered neuroprotective, as neurons can utilize astrocytic GSTM2 for efficient detoxification of neuronal aminochrome, enhancing resilience to oxidative stress. Dashed arrows indicate putative or less well-established pathways. DA, dopamine; DAT, DA transporter; PMAT, plasma membrane monoamine transporter; NET, Norepinephrine transporter; OCT3, organic cation transporter 3; VMAT2, vesicular monoamine transporter 2; MAO, monoamine oxidase; COMT, catechol-O-methyltransferase; GSTM2, glutathione S-transferase mu 2. Illustration based on work from Segura-Aguilar and coworkers (*26, 53*) and further extended in this work. Created in BioRender.

In sum, our work using human iPSC-derived models demonstrates that DJ-1 loss in astrocytes impairs both mitochondrial and DA metabolic adaptability, disrupting key glial defences against dopaminergic stress. Recognition of these astrocytic vulnerabilities should inform the development of broader therapeutic strategies for PD that target both neuronal and glial contributors.

## Materials and Methods

### Subjects

Included in this study are iPSCs that were already generated from fibroblasts from a healthy control individual (Ctrl) and its isogenic line carrying a CRISPR-Cas9 inserted mutation of c.192G>C in the *PARK7* gene (DJ-1 mut), both previously described and characterized as “C4” and “C4Mut”, respectively (*55*). The study was approved by the ethics committee of the LMU Munich. The participant gave written and informed consent.

### Culture of human iPSCs

iPSCs were grown in mTeSR (Stemcell Technologies, #100-0276) on Cultrex-coated (R&D systems, #3434-010-02) plates and passaged manually every 5 to 7 days. All lines were routinely tested for mycoplasma contamination.

### Astrocyte differentiation

Astrocyte differentiation was carried out from iPSCs via small molecule-derived neural precursor cells (smNPC) and human neural stem cells (hNSC), adapted from previously described protocols (*56*) (*38*). From day 1 to 4 of differentiation, iPSCs were exposed to N2B27 medium containing 50% DMEM/F12 w/o HEPES (Thermo Fisher, #21331046), 50% NeuroBasal medium (Thermo Fisher, #21103049), 0.5% N2 supplement (Life/Tech, #17502048), 1% B27 supplement lacking vitamin A (Life/Tech, #12587-010), 1 % pen/strep (Life/Tech, #15140-163) and 1 % GlutaMAX (Life/Tech, #35050-061) with 10μM SB-431542 (R&D Systems, #1614), 1μM dorsomorphin (Sigma, #P5499), 3μM CHIR 99021 (Stemgent, #04-0004-10) and 0.5μM PMA (Merck, #540220). On day five, media was changed to smNPC medium, consisting of N2B27 with 3μM CHIR 99021, 0.5μM PMA and 150μM ascorbic acid (Sigma, #A8960). On day 7, neuroepithelial structures were manually picked with a P200 pipette, dissociated and plated onto a Cultrex-coated 12-well plate. smNPCs were passaged 1:20 using Accutase (Sigma Aldrich, #A6964) every 7 days. After obtaining pure smNPC cultures, 4×10^^5^ smNPCs were seeded per well of a 6-well plate to start hNSC differentiation. After 2 days, smNPC medium was supplemented with 20ng/ml FGF-2 (Peprotech, #100-18B). On day 4, cells were split into hNSC medium consisting of DMEM/F12 w/o HEPES, 1% N2 supplement, 2% B27 supplement with vitamin A, GlutaMAX, penicillin/streptomycin and 40ng/ml EGF (Peprotech, #AF-100-15-1mg), 4ng/ml FGF-2 and 1.5ng/ml LIF (Peprotech, #AF-300-05). To start the astrocyte differentiation, hNSCs were plated into Cultrex-coated T25 flasks in hNSC medium. After two days, medium was changed to astrocyte medium, consisting of DMEM/F12 w/o HEPES with 1 % penicillin/streptomycin, 1 % GlutaMAX and 1 % fetal bovine serum (Life/Tech, #10270-106). At day 40 of differentiation cells were passaged 1:1 to remove any possible contamination with post-mitotic neurons. Astrocytes were maintained in the flask until mature at day 70 of differentiation, after which they were passaged or harvested for subsequent experiments.

### Dopaminergic neuron differentiation

Differentiation of human iPSCs into dopaminergic neurons followed previously published protocols (*57*) and was done as previously described (*5*). Briefly, iPSCs were dissociated by Accutase into single cells and plated at a density of 6×10^^5^ per well of a 6-well plate. Once the cells were >95% confluent, differentiation medium (Knockout DMEM, Knockout Serum Replacement (KOSR), 1% penicillin-streptomycin, 1% L-glutamine, 1% NEAA and 1mM beta-mercaptoethanol) was added. From day 5 to 10 of differentiation the medium was gradually weaned onto Neurobasal media supplemented with Neurocult SM1, 1% penicillin-streptomycin, and 1% L-glutamine. On day 12 after neuronal induction, cells were passaged manually by cutting 1-2mm chunks and plating them onto poly-D-lysine/laminin (Sigma Aldrich, #P1149 and #11243217001) coated 10-cm dishes. Reaching day 30 of differentiation, these cells were detached and dissociated into single cells using Accutase and plated into poly-D-lysine/laminin-coated 12-well plates. Neuronal differentiation factors were removed at day 40 of differentiation and cells were maintained in Neurobasal media at day 70 of differentiation when they reached maturity.

### Mass spectrometry

Cells were harvested for mass spectrometry by incubation in 1mM EDTA for 10 min and centrifuged at 300g for 10min. Cells were lysed in STET lysis buffer supplemented with protease inhibitor (Sigma-Aldrich, #539133) and the protein concentration was determined with a BCA assay. A protein amount of 20 µg per sample was subjected to digestion using the single-pot, solid-phase-enhanced sample preparation (SP3) method (*58*) with some modifications. DNA and RNA were digested using 25 units of Benzonase (Sigma-Aldrich) at 37°C for 30 min. Afterwards proteins were reduced with 15 mM dithiothreitol for 30 min at 37°C followed by alkylation of cysteine residues with 60 mM iodoacetamide for 30 min at room temperature. Proteins were bound to 40 µg of a 1:1 mixture of hydrophobic and hydrophilic magnetic carboxylate-modified Sera-Mag SpeedBeads (Cytiva) for 30 min in 70% (v/v) acetonitrile. Beads were washed four times with 80% (v/v) ethanol. Protein digestion was performed with 0.25 µg LysC (Promega; incubation 1 h at 37°C) and trypsin (Promega; incubation 16h at room temperature). Peptides were collected, filtered with 0.22 µm Corning Spin-X centrifuge tube filters (Sigma-Aldrich) and dried by vacuum centrifugation. Peptides were dissolved in 0.1% (v/v) formic acid (FA) and the concentration was estimated using the Qubit protein assay (Thermo Fisher).

A peptide amount of 300 ng per sample was analysed on a nano-Elute nanoHPLC coupled online to a timsTOF pro mass spectrometer (Bruker). Samples were separated on a self-packed C18 column (15 cm × 75 µm; ReproSil-Pur 120 C18-AQ resin, Dr. Maisch GmbH) using a binary gradient of water and acetonitrile supplemented with 0.1% (v/v) FA (0 minutes, 2% B; 2 minutes, 5% B; 62 minutes, 24% B; 72 minutes, 35% B; 75 minutes, 60% B). A Data Independent Acquisition Parallel Accumulation– Serial Fragmentation (diaPASEF) method was used. A ramp time of 100 ms was used for the trapped ion mobility spectrometry (TIMS). Each scan cycle included one TIMS full MS scan and 13 PASEF scans of 2 windows with a width of 27 m/z covering a m/z range of 350-1001 m/z.Protein label-free quantification (LFQ) data analysis was performed using the software DIA-NN version 2.2 (*59*). The raw data was searched against a one protein per gene database from species (UniProt, 20662 entries, download: 2024-12-09) and a database with potential contaminants (233 entries) using a library free search. Trypsin was defined as protease and two missed cleavages were allowed. Oxidation of methionines and acetylation of protein N-termini were defined as variable modifications, whereas carbamidomethylation of cysteines was defined as fixed modification. Variable modifications were restricted to two per peptide. The precursor and fragment ion m/z ranges were limited from 350 to 1001 and 200 to 1700, respectively. Precursor charge states were set to 2-4 charges. An FDR threshold of 1% was applied for peptide and protein identifications. Mass accuracy thresholds and ion mobility windows were automatically adjusted by the software. The match between runs and RT-dependent cross-run normalization options were enabled.

The software Perseus version 1.6.2.3 (*60*) was used for further data analysis. Protein LFQ data was filtered for identifications with at least two peptides. Protein LFQ intensities were log2 transformed. Changes in protein abundances between the different groups were calculated and a two-sided Student’s t-test was applied comparing the log2 transformed LFQ intensities between the groups. At least three valid values per group were required for statistical testing. To account for multiple hypotheses, a permutation-based FDR correction (*61*) was applied (p = 0.05; s_0_ = 0.1).

### Measurement of oxygen consumption rate

Mitochondrial oxygen consumption rate (OCR) was measured using the Seahorse XF Cell Mito Stress Test kit (Agilent, #103015-100) on a Seahorse Analyzer XFe96 (Agilent). One day prior to analysis, astrocytes were seeded at a density of 5×10^^4^ on a Cultrex-coated 96-well plate and the cartridge was hydrated overnight in a CO_2_-free incubator at 37°C. On the day of the experiment, cell media was replaced with assay medium (Seahorse XF DMEM, 2mM L-Glutamine, 1mM Sodium Pyruvate, 1g/L Glucose (Agilent, # 103680-100)) and plates were preincubated in a CO_2_-free incubator at 37°C for 1h to equilibrate. XF Cell stress test compounds were diluted according to the manufacturer’s recommendations and loaded into the cartridge, after which the assay was started. The assay followed standard protocol, with initial baseline measurements, after which oligomycin (2μM), FCCP (1μM), and rotenone/antimycin A (0.5μM) were sequentially injected, with three measurement cycles each. For normalization of cell number, DAPI stained nuclei were counted using the BioTek Cytation 1 Cell Imaging Multimode Reader. Data was analysed via the Seahorse Wave software and GraphPad Prism.

### Dopamine treatment

Astrocytes at day 70 of differentiation were plated at a density of 6×10^^5^ in 6-well plates and treated with 60µM dopamine hydrochloride (Sigma Aldrich, #H8502-5G) or PBS in astrocyte media for 5 days, with media being refreshed every day. Cells were harvested on day 5 of treatment for consecutive analysis.

### High-performance liquid chromatography (HPLC)

Intracellular catecholamine content was measured with reverse phase high-performance liquid chromatography (HPLC) using a hyperchrome-hplc-column (Prontosil, #120-3-CL8-AQ) and an electrochemical detector. After DA treatment, astrocytes were harvested in PBS, centrifuged 10min, 500g at 4°C and resuspended in 100µl of 1µM perchloric acid (Roth, #9216) containing 300nM 3,4 dihydroxybenzylamine (DHBA, Sigma, #858781). Cell lysates were sonicated and centrifuged at 17,000g for 10 min at 4°C. Supernatants were filtered through 0.45 μm PTFE filter vials (Thomson, #TI35540) and 20µl per sample was injected on the HPLC for analysis of dopamine levels and metabolites. Quantification of dopamine and metabolites was done by comparing the peak areas of a known amount of standard. Concentration of total protein measured by Pierce BVA Protein Assay Kit (BCA, Thermo Scientific, #23225) was used for normalization.

### Protein extraction and western blot analysis

Cells were harvested by scraping in ice-cold PBS (neurons) or detaching via Accutase (astrocytes) and centrifuged at 300g for 5 min. Pellets were resuspended in lysis buffer (containing 1% Triton X-100, 10% glycerol, 150mM NaCl, 25mM Hepes pH7.4, 1mM EDTA, 1.5mM MgCl2, and proteinase inhibitor cocktail) and homogenized before being incubated on ice for 20 min and centrifuged at 100,000g for 30 min at 4°C. The protein concentration in the supernatant was determined using the bicinchoninic acid (BCA) assay. For immunoblotting, 8µg protein were run on an SDS-PAGE 4-20% gel (Thermo, #XP04200BOX) and transferred to a PVDF membrane (Merck, #IPFL00010). After blocking with Intercept blocking reagent (Licor, #927-70003) for 1h, the following primary antibodies were used for incubation o/n at 4°C: DAT (Santa Cruz, #sc-32259, 1:500), DJ-1 (abcam, #ab76008, 1:1000); GAPDH (Millipore, #MAB374, 1:10000); GFAP (Millipore, #MAB360, 1:1000); GSTM-2 (Santa Cruz, #sc-376486, 1:500); MAP2 (Sigma, #M4403, 1:1000); NQO1 (abcam, #ab80588, 1:500). Blots were probed with IRDye fluorescent secondary antibodies (Li-Cor, # 926-68070, # 926-32211) for 1h and imaged using the Odyssey XF Imaging System (Li-Cor).

### Detection of dopamine quinone species by near infrared fluorescence (niRF) assay

After cell lysis, the Triton insoluble pellet was further processed in order to measure oxidized dopamine content. First, the pellets were further extracted in 2% SDS/50mM Tris pH7.4, in a volume that was normalized according to the protein content of the T-soluble fraction. Samples were boiled at 95°C for 10 min and sonicated, before being centrifuged at 150,000g for 30 min. The insoluble pellets were further extracted by incubation in 1N NaOH overnight at 55°C in half the volume of the SDS-buffer volume.

Samples were sonicated again and lyophilized in a Speed-Vac Concentrator until dry, before being washed once in H_2_O and lyophilized again. Then, samples were taken up in H_2_O, at the same volume as NaOH, and 5ul of the solution were dropped onto a Biodyne Transfer Membrane (Pall, #-60208). Membranes were imaged using the 700 channel of the Odyssee XF Imaging System and spot intensities were measured for quantification.

### Immunofluorescence staining and image analysis

For immunofluorescence analysis, cells grown on coverslips were fixed in 4% formaldehyde for 15 min and permeabilised and blocked in PBS, 0.4% Triton-X, 10% donkey serum and 2% BSA for 1h at RT. Cells were incubated with primary antibodies in in PBS, 0.1% Triton-X, 1% donkey serum and 0.2% BSA over night at 4°C, using the following antibodies: GFAP (Synaptic Systems #173004 1:500), S100ß (Abcam,# ab52642 1:250), Vimentin (Cell Signaling, #5741S 1:250), Sox9 (Merck, #5535 1:500), EAAT2 (Santa Cruz, 365634 1:100), MAP2 (Sigma, #M4403 1:250). After washing, coverslips were incubated with anti-mouse or anti-rabbit secondary antibodies (Invitrogen #A11036; 1:500; Invitrogen #A11029; 1:500) for one hour and mounted in Hoechst-containing mounting media (Thermo Fischer, #P36981). Imaging was carried out using a Leica TCS SP5 confocal microscope (Leica Microsystems).

### Flow Cytometry

Astrocytes were detached with Accutase, centrifuged at 300g for 5 min and resuspended in PBS. Cells were fixed with 4% PFA for 15 min and washed with PBS/5% BSA, before being incubated in Saponin buffer (0.1% Saponin, 5% BSA, PBS) for 20min at 4°C. Afterwards, cells were pelleted by centrifugation at 700g for 5min and resuspended in 100µl primary antibody in saponin buffer. After 1hr incubation at 4°C, cells were washed 3 times with PBS/5%BSA and incubated in secondary antibody solution in PBS/5% BSA for 30min at 4°C. After two more washes with PBS/5% BSA, cells were resuspended in 250µl PBS and analysed using the BD LSRFortessa Cell Analyser. FlowJo software was used to measure the mean fluorescence intensity of the astrocytes.

### Glutamate Assay

Astrocytes at day 70 of differentiation were plated at a density of 6×10^^5^ in 6-well plates and treated with 100µM L-glutamic acid or PBS as a vehicle (Sigma, #G8415) for 4 hours. Cells were washed, harvested and uptake was measured using a glutamate assay kit (Abcam, #ab83389). Values were normalised to protein concentration determined by BCA assay (Thermo, #23225)

## Statistical analysis

Comparison between two groups were performed using Student’s t-test, whereas comparison of multiple groups and conditions was performed using two-way ANOVA followed by post-hoc Tukey test. Statistical analyses were performed in Perseus (volcano plot) and GraphPad Prism. P-values less than 0.05 were considered significant. In Figure 3 and 4, all blots are indicated as fold change relative to the control condition and loading control. All errors bars shown in the figures are standard error of the mean (SEM).

## Acknowledgements

This work was supported by the Deutsche Forschungsgemeinschaft (DFG, German Research Foundation) under the Heisenberg Programme (Project No. 447395247) (to L.F.B.) and under Germany’s Excellence Strategy within the framework of the Munich Cluster of Systems Neurology (EXC 2145 SyNergy—ID 390857198) (to L.F.B. and S.F.L.), and by a doctoral scholarship from the Hans and Ilse Breuer Foundation (to A.W.). The project was also supported through the Bundesministerium für Forschung, Technologie und Raumfahrt (BMFTR, FKZ: 01ED2402A) under the aegis of JPND (to S.F.L.). The work of R.K. was supported by grants within the National Centre of Excellence in Research on Parkinson’s Disease (NCER-PD) funded by the Luxembourg National Research Fund (FNR/NCER13/BM/11264123); the PEARL program of the FNR (PEARL/P13/6682797); and a CORE grant of the FNR (Luxembourg National Research Fund (C17/BM/11676395). The work of R.K and I.B. is supported by European Commission’s Horizon Europe Programme under grant no. 101080249. We thank Christian Haass for providing equipment and assistance.

## Data and materials availability

All data needed to evaluate the conclusions in the paper are present in the paper and/or the Supplementary Materials.

## Supplemental Information

**Supplementary Figure 1.**
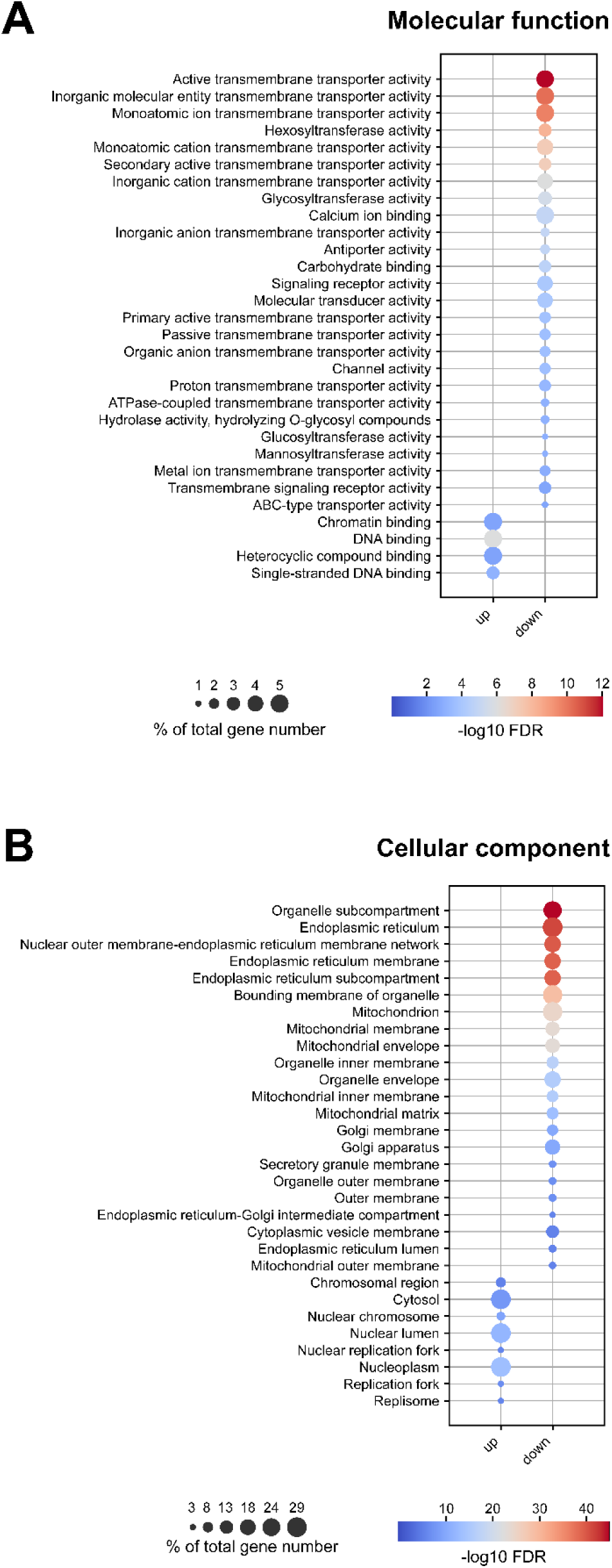
Gene ontology enrichment analyses for molecular function and cellular component of control and DJ-1-deficient astrocytes. (**A-B**) Gene ontology (GO) enrichment analyses for (A) molecular function (GOTERM_MF_DIRECT) and (B) cellular component (GOTERM_CC_DIRECT) were performed using DAVID (v 6.8) (*62*). Differentially expressed proteins (T-test p-value < 0.05 and log2 fold change > 0.5 or < −0.5) were compared to the background of all proteins detected under both conditions. In the dot plots, dot size represents the total gene number of a term in percent, while colour indicates the -log10-transformed FDR. The top 30 terms for either enriched (up) or downregulated (down) proteins are shown.

